# Nuclear envelope assembly defects link mitotic errors to chromothripsis

**DOI:** 10.1101/263392

**Authors:** Shiwei Liu, Mijung Kwon, Mark Mannino, Nachen Yang, Alexey Khodjakov, David Pellman

## Abstract

Defects in the architecture or integrity of the nuclear envelope (NE) are associated with a variety of human diseases^1^. Micronuclei, one common nuclear aberration, are an origin for chromothripsis^2,3^, a catastrophic mutational process commonly observed in cancer genomes and other contexts^4-6^. Micronuclei have a defective NE, with the extensive chromosome fragmentation that generates chromothripsis occurring after abrupt, spontaneous loss of NE integrity^7^. After NE disruption, the exposed cytoplasmic DNA can additionally initiate proinflammatory signaling linked to senescence, metastasis, and the immune clearance of tumor cells^8^. Despite its broad physiological impact, the basis for the nuclear envelope fragility of micronuclei is unknown. Here we demonstrate that micronuclei undergo markedly defective NE assembly: Only “core” NE proteins^9,10^ assemble efficiently on lagging chromosomes whereas “non-core” NE proteins^9,10^, including nuclear pore complexes (NPCs), fail to properly assemble. Consequently, micronuclei have impaired nuclear import, and key nuclear proteins required to maintain the integrity of the NE and the genome fail to accumulate normally. We show that densely bundled spindle microtubules inhibit non-core NE assembly, leading to an irreversible NE assembly defect. Accordingly, experimental manipulations that position missegregated chromosomes away from the spindle correct defective NE assembly, prevent spontaneous NE disruption, and suppress DNA damage in micronuclei. Our findings indicate that chromosome segregation and NE assembly are only loosely coordinated through the timing of mitotic spindle disassembly. The absence of precise regulatory controls can explain why errors during mitotic exit are frequent, and a major trigger for catastrophic genome rearrangements^5,6^.

---

During normal mitotic exit, NE proteins transiently form two domains around decondensing chromosomes^1,9,10^. The group of “core” NE proteins, which include the membrane protein emerin, and BAF (barrier-to-autointegration factor), concentrate on the chromosome mass adjacent to central spindle or spindle pole microtubules; the “non-core” group of proteins, which include nuclear pore complex proteins (NPCs) and the Lamin B receptor (LBR), assemble on the chromosome periphery away from the spindle (Extended Data Fig. 1a)^9,10^. After mitotic exit, these domains become intermingled, with fragments of the core domain persisting as “pore-free” islands that are then slowly populated by NPCs during interphase^11,12^.

**Figure 1.**
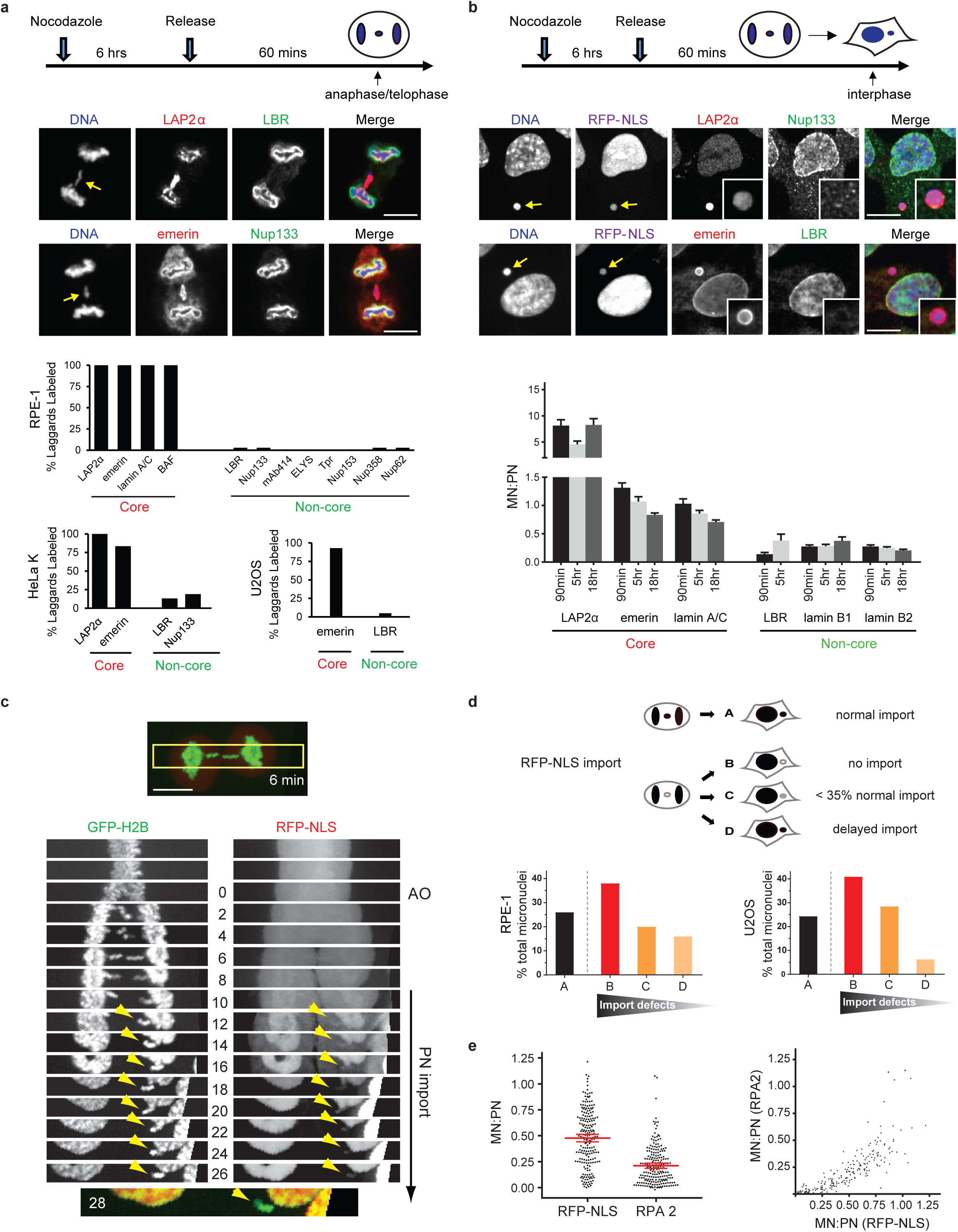
Micronuclei undergo defective NE assembly. **a, b.** Defective non-core NE protein recruitment to lagging chromosomes (**a**) or micronuclei (MN, **b**). Top panels: Experimental scheme. Middle panels: Representative images of RPE-1 cells with lagging chromosomes or MN (arrows, insets for enlarged images). Red letters: core NE proteins; Green: non-core proteins. (**a**) Bottom: quantification of the results (n > 50; ≥ 2 experiments). (**b**) Bottom: The fluorescence intensity (FI) ratio of the indicated proteins in intact MN relative to primary nucleus (PN) in RPE-1 cells at indicated timepoints after release from nocodazole block (mean with 95% CI, n > 100; 2 experiments). mAb414 detects nucleoporins. **c, d**, Impaired nuclear import in MN. **c,** Kymograph of RPE-1 cell after synchronization as in **a**, 2 min intervals (t=0 is anaphase onset, AO). Kymograph is from the boxed region of top image. Merged image at bottom shows poor accumulation of the import reporter, RFP fused to a nuclear localization signal (RFP-NLS) in the MN, which persisted for at least 2h (not shown). Arrowheads: newly formed MN. **d,** Top: Cartoon depicting patterns of import to MN: (A) Normal import kinetics; (B) No detectable import; (C) Delayed import with a persistent defect (up to 35% of PN) and (D) Delayed import with eventual normal RFP-NLS accumulation. Bottom: Percentage of cells corresponding to the categories above in the indicated cell lines (n=21 for RPE-1 and n=17 for U2OS). **e**, Left: The MN/PN FI ratio of RPA2 and RFP-NLS in RPE-1 cells ∼1h post AO (mean with 95% CI, n > 200, 2 experiments). Right: Defective accumulation of RPA2 and RFP-NLS are correlated (r = 0.9039, P < 0.0001, Spearman’s correlation analysis). Scale bars, 10 μm.

To understand the basis for NE abnormalities of micronuclei, we asked whether postmitotic NE assembly on lagging chromosomes occurs in this same spatiotemporal pattern. In three cell lines, lagging chromosomes were generated in synchronized cells by recovery from nocodazole-induced mitotic arrest or by short-term inhibition of the spindle assembly checkpoint with a MPS1 kinase inhibitor. Immunofluorescence staining or live-cell imaging of telophase cells revealed that core proteins were recruited to lagging chromosomes at equivalent or higher levels than observed on the main chromosome mass. By contrast, the non-core proteins were strikingly depleted from lagging chromosomes (Fig. 1a, Extended Data Fig. 1b-d). Likewise, chromatin bridges formed after nocodazole release or after partial depletion of the condensin SMC2 displayed the same core-only NE protein composition (Extended Data Fig. 1e, f).Thus, only a subset of NE proteins assembles on chromosomes that lag within the region of the central spindle during telophase.

A similar reduction of non-core NE proteins was observed on micronuclei formed from lagging chromosomes (Fig. 1b, Extended Data Fig. 2a-c). Consistent with the reduced assembly of NPCs on micronuclei, high temporal resolution live-cell imaging of cells demonstrated significant nuclear import defects in micronuclei, with some variation in the extent of the defect (two import reporters in two cell lines, Fig. 1c, d, Extended Data Fig. 3a, b, Supplementary Videos 1 and 2). Micronuclei fail to normally accumulate key nuclear proteins, including replication protein A (RPA), a key dosage-sensitive regulator of DNA replication and repair^13^, and B-type lamins, known to be required to maintain NE integrity (Fig. 1b, e, Extended Data Figs. 2a-c, 3c, 3d)^7,14^. For individual cells, the extent of the defect in accumulating RPA and a general import reporter were strongly correlated, suggesting that micronuclei have a global import defect rather than selective import pathway defects (Fig.1e). Thus, reconciling differing reports from prior literature^2,7,15-18^, these data indicate that micronuclei undergo aberrant NE and NPC assembly, leading to defective import and accumulation of many proteins, including those required for genome stability and NE integrity.

**Figure 2.**
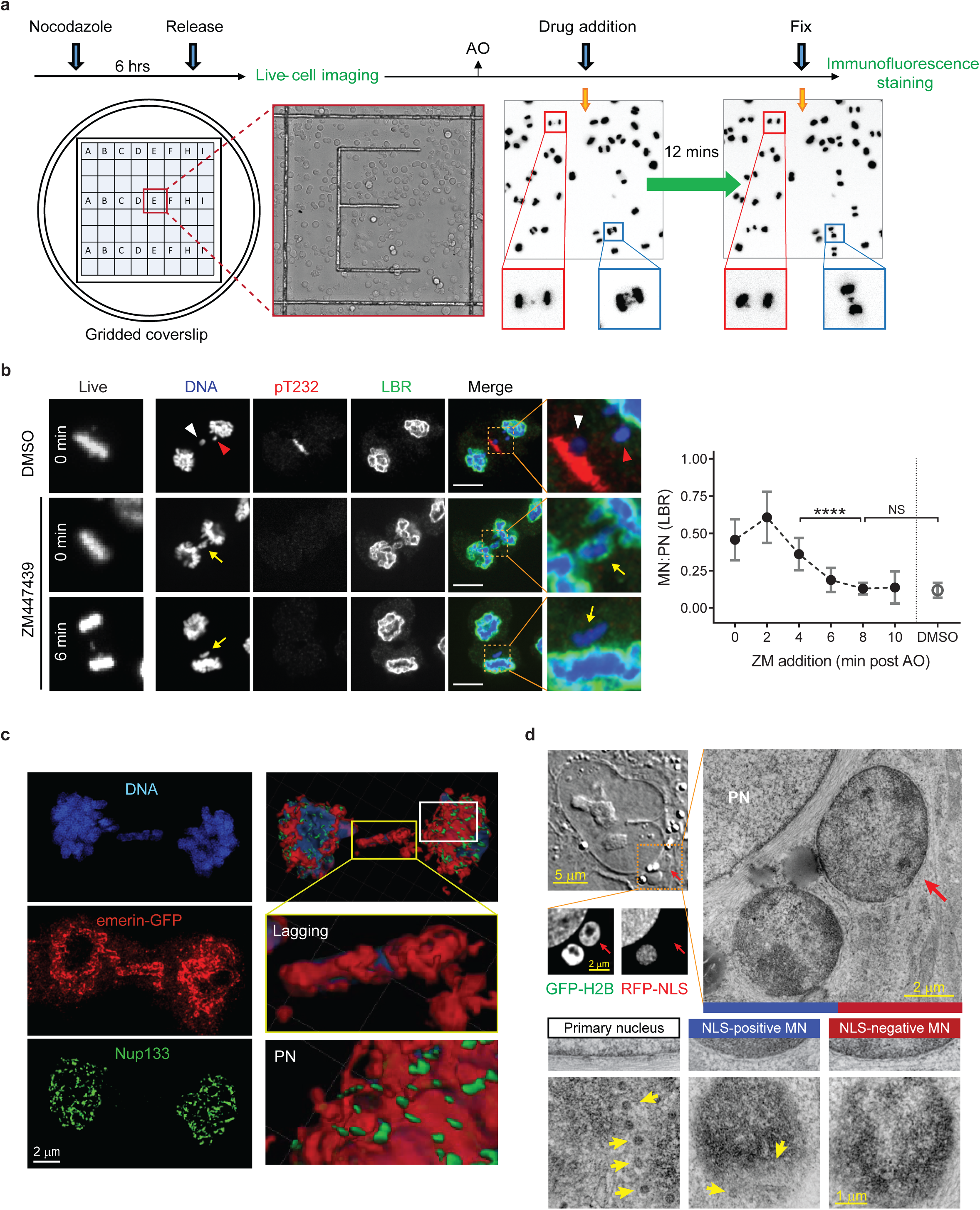
NE assembly defect of lagging chromosomes becomes irreversible in telophase. **a,** Scheme of the live-cell/ fixed cell imaging protocol. RFP-H2B-expressing RPE-1 cells were plated on gridded dishes to identify cells of interest. Cells were imaged at 2 min intervals during the experiments. Images of two live cells (red and blue boxes) upon (left) or after (right, prior to fixation) ZM447439 addition. **b**, Aurora B inhibition after late-anaphase fails to restore non-core (LBR) assembly to lagging chromosomes (arrows). Left column: representative cells (0 or 6 min post AO) at the time of ZM447439 (ZM) or DMSO addition. Right columns: Cells labeled for DNA (blue), phospho-T232 Aurora B (red) and LBR (green). For the control (DMSO) samples, lagging chromosomes fail to recruit LBR whether they are proximal (white arrowhead) or distal (red arrowhead) to activated Aurora B (pT232). Scale bars, 10 μm. Graph on right: MN/PN LBR FI ratio in cells exposed to ZM at the indicated times (mean with 95% CI, n > 50 each timepoint, ≥ 3 experiments, for simplicity, lagging chromosomes are designated as “MN”). **** P < 0.0001, NS: not significant, Mann-Whitney test. **c,** Near-continuous assembly of core membrane protein around the lagging chromosome. Images of 3D-SIM (structured illumination microscopy, left) and Imaris surface renderings (right) of an emerin-GFP-expressing RPE-1 cell. Enlarged images show lagging chromosomes (yellow box) and PN (white box): DNA (blue), emerin (red), Nup133 (green) (representative of 22 cells). **d,** Correlative light and electron microscopy (CLEM) showing membranous NE (enlarged images in middle panels, cross sections) but reduced NPCs (yellow arrows in bottom panels, tangential sections) on intact (RFP-NLS-positive) and newly disrupted MN (RPF-NLS negative, red arrows). Left, top: DIC image of an RPE-1 cell just after loss of NE integrity of one MN. Right, top: EM image; fixation was < 20 min after MN disruption. Enlarged images (middle and bottom panels) are at the same magnification.

**Figure 3.**
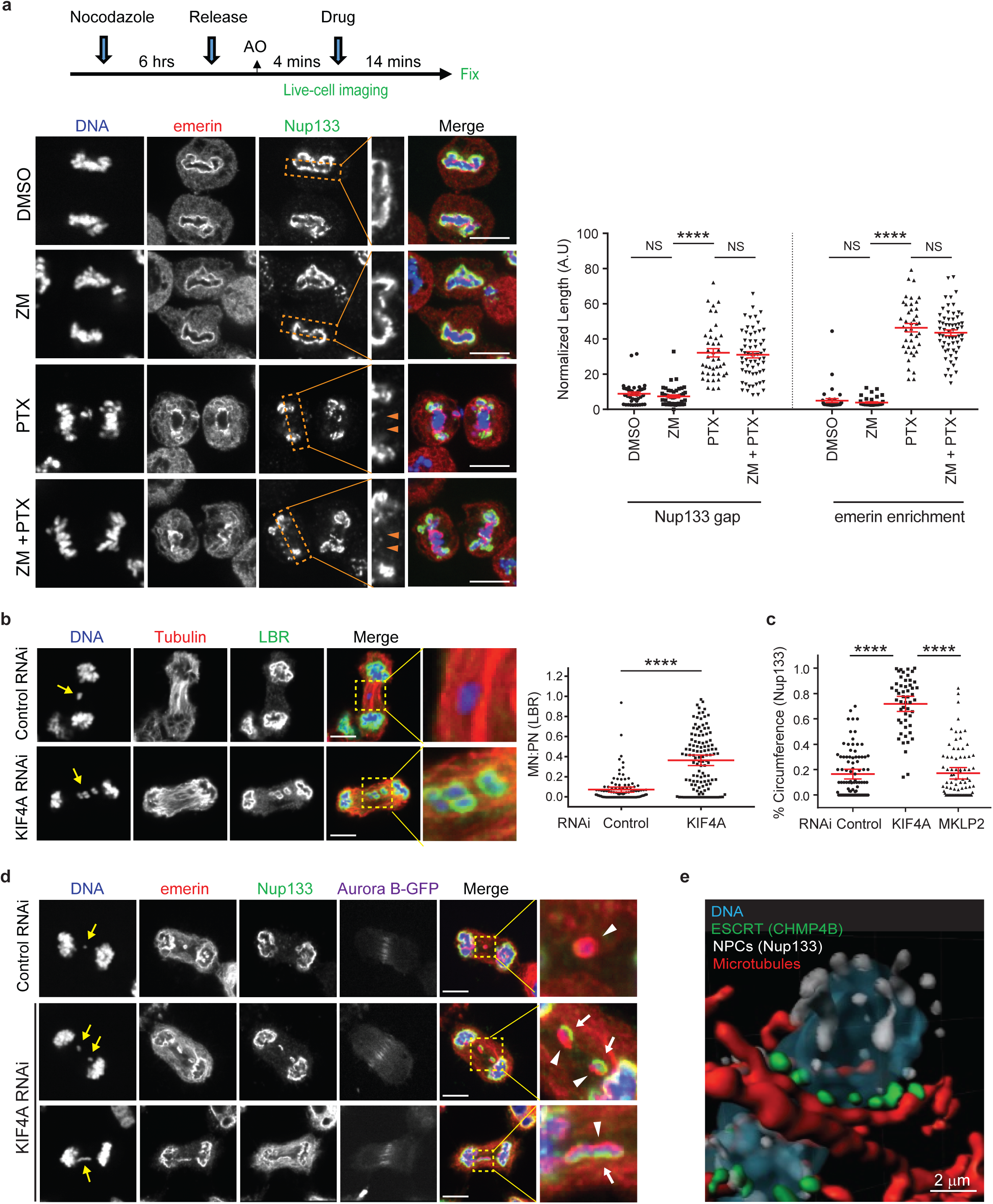
Independent of Aurora B, bundled microtubules inhibit non-core NE recruitment to lagging chromosomes. **a,** Paclitaxel (PTX) inhibits non-core assembly. Left, top: Experimental scheme. Left, bottom: representative images of cells after drug treatments. Merged image: DNA (blue), emerin (red) and Nup133 (green). Enlarged images from the boxed regions show the non-core (Nup133) gap (orange arrowheads). Right: quantification of the results (mean with 95% CI, n > 40, 2 experiments). **** P < 0.0001, NS: not significant, Mann-Whitney test. **b**, Restoration of LBR to some lagging chromosomes after KIF4A depletion (RPE-1 cells). Synchronization as in **Fig. 1a**. Left: Representative images: DNA (blue), tubulin (red) and LBR (green). Right: MN/PN LBR FI ratio (mean with 95% CI, n > 100, 3 experiments). **** P < 0.0001, Mann-Whitney test. **c, d**, Small-scale core/non-core domain separation on lagging chromosomes after KIF4A depletion. Synchronization as in **Extended Data Fig. 1c**. **c,** The percentage of the lagging chromosome circumference with Nup133 in HeLa K cells (mean with 95% CI, n > 60, 2 experiments). *** P < 0.0001, Mann-Whitney test. **d,** Representative images of Aurora B-GFP-expressing HeLa K cells. Merged and enlarged image: emerin (red, white arrowheads) Nup133 (green, white arrows). Scale bars, 10 μm. **e,** Imaris surface three-dimensional renderings from SIM image showing recruitment of Nup133 (white) to the region of a lagging chromosome depleted of CHMP4B (green) and microtubules (red) from a KIF4A-depleted HeLa K cell (representative of 4 lagging chromosomes).

We considered the possibility that the non-core recruitment defect on lagging chromosomes could be explained by a previously proposed “chromosome separation checkpoint”^17^. Under this model, the position of lagging chromosomes is monitored throughout mitotic exit by the spindle midzone-centered Aurora B phosphorylation gradient^19^, postulated to block NE assembly until membrane-free, lagging chromosomes can be incorporated into the main chromosome mass, ensuring the formation of a single nucleus. However, several observations were inconsistent with this model. First, core membrane proteins, which were not previously examined^17^, assemble on lagging chromosomes and chromosome bridges^20^, independent of their position relative to the spindle midzone (Extended Data Fig. 4a). Thus, an NPC-depleted NE does form prior to the completion of chromosome segregation. Second, chromosome bridges are uniformly depleted for non-core proteins, showing no obvious gradient (Extended Data Figs. 1e, f, 4a). Finally, we inhibited Aurora B transport to the spindle midzone by siRNA-mediated knockdown of the kinesin MKLP2^21^. After MKLP2 knockdown, cells lacking detectable midzone Aurora B nevertheless failed to recruit non-core proteins to lagging chromosomes (Extended Data Fig. 4b), indicating that the gradient is not required for inhibition of non-core NE assembly.

**Figure 4.**
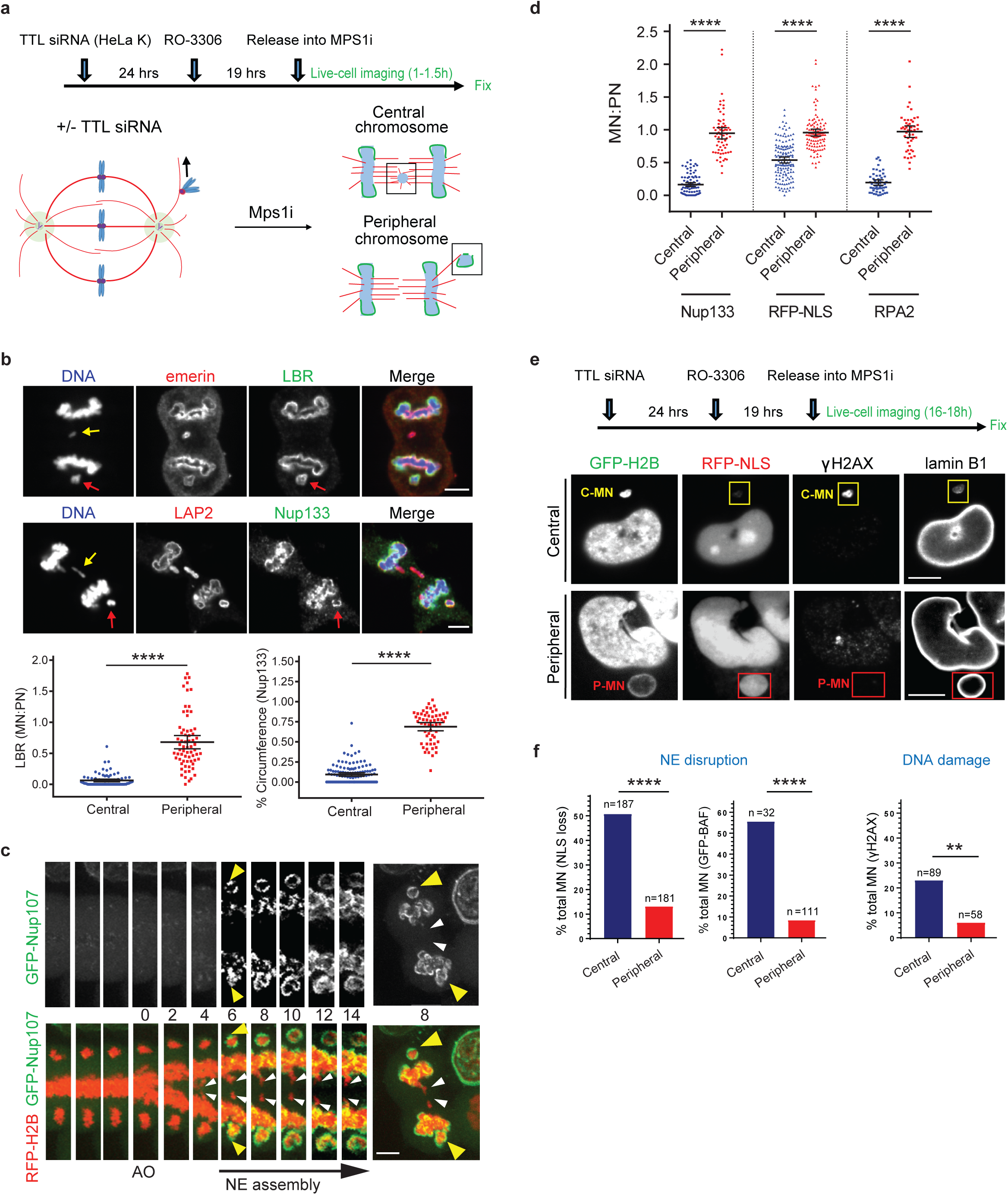
Peripheral localization of missegregated chromosomes corrects defects of micronuclei. **a**, Experimental scheme to generate missegregated chromosomes (MPS1i with or without TTL knockdown) positioned within or away from the spindle (microtubules: red; non-core NE: green). Cells were imaged at 4 min intervals throughout mitosis. **b, c**, Non-core NE assembles on peripheral chromosomes but not to central chromosomes. **b,** Top: Representative images of RPE-1 cells with central (yellow arrows) and peripheral (red arrows) chromosomes labeled with core (red) or non-core (green) proteins. Bottom: Quantification as in **Fig. 3b, c**(mean with 95% CI, n > 50, 2 experiments). ^****^ P < 0.0001, Mann-Whitney test. **c,** Similar result as in **b** from live-cell imaging of HeLa K cell expressing RFP-H2B (red) and GFP-Nup107 (green) (Supplementary Video 4). AO is t=0 (min). **d-f,** Restoration of function for MN from peripheral chromosomes. **d,** FI ratios for the indicated proteins, comparing MN from peripheral with central chromosomes (RPE-1 cells, scheme as in **a**) (mean with 95% CI, n > 50, 2 experiments). ^****^ P < 0.0001, Mann-Whitney test. **e,** Top: Experimental scheme. Bottom: Representative images of HeLa K cells (C-MN from central chromosomes; P-MN from peripheral chromosomes). **f,** Graphs of the results. Left: NE integrity monitored by loss of NLS-RFP (5 experiments). Middle: NE integrity monitored by hyper-accumulation of GFP-BAF (3 experiments). Right: DNA damage (3 experiments). ** P < 0.05, ^****^ P < 0.0001, Fisher’s exact Test. Scale bars, 5 μm.

We designed an experiment to directly test the main prediction of the checkpoint model that the position of lagging chromosomes is monitored continuously throughout mitotic exit by Aurora B. Cells were released from a mitotic arrest, followed by live-cell imaging at high temporal resolution, treated with an Aurora B inhibitor (ZM447439), and then fixed and labeled to assess NE assembly (Fig. 2a). Because cells release from the mitotic block with only partial synchrony, this approach enabled us to identify cells exposed to Aurora B inhibition at all intervals from metaphase through telophase. Strikingly, we found that Aurora B inhibition restored non-core protein assembly to lagging chromosomes only if it occurred early, at or shortly after anaphase onset. However, there was no effect if Aurora B inhibition occurred later, ~6-8 min after anaphase onset (Fig. 2b). Thus, global Aurora B activity inhibits NE assembly up until early anaphase, but has no effect during the critical time period where a chromosome separation checkpoint would need to operate. Instead, the lagging chromosomes appear to become irreversibly defective for non-core NE recruitment in telophase. This irreversible effect is unlikely to be caused by persistent, Aurora B-mediated association of condensin I with lagging chromosomes^17^ because siRNA-mediated knockdown of SMC2 did not rescue non-core NE assembly (Extended Data Fig. 4c).

The formation of a nearly continuously enclosed, core-only NE on the lagging chromosome could explain the irreversible defect in the recruitment of NPCs and other non-core proteins. This idea is consistent with the observation that *Xenopus* egg extracts lacking the Nup107-160 complex assemble NPCs only if the purified complex is added back prior to the formation of a continuous NPC-free NE^22^. Importantly, telophase lagging chromosomes are already ensheathed by an apparently continuous layer of the core-membrane protein emerin (Fig. 2c). Concomitantly, ESCRT-III components, which are thought to seal small membrane gaps during NE assembly^23,24^, associate and dissociate from lagging chromosomes approximately on schedule (Extended Data Fig. 5a, b, Supplementary Video 3). These ESCRT-III kinetics suggest that the micronuclear membrane likely undergoes significant membrane fusion. Accordingly, correlated light and electron microscopy (CLEM) demonstrated that the NPC-deficient NE formed around lagging chromosomes persists on micronuclei throughout interphase (Fig. 2d). As expected for defective NPC assembly and therefore defective import, chromatin within these structures was hyper-condensed (Fig. 2d)^25^.

Although our experiments argue against spatial regulation of NE assembly by the Aurora B gradient (above and Extended Data Fig. 4a, b), we are still left with the question of why global inhibition of Aurora B in early anaphase allows non-core NE assembly on lagging chromosomes (Fig. 2b). We considered the possibility that this effect on NE assembly could be an indirect consequence of Aurora B’s well-characterized role in stabilizing spindle microtubules, which increases both the mass and bundling of spindle microtubules (Extended Data Fig. 6a)^26^. Supporting the idea that microtubules adjacent to chromosomes can prevent non-core NE assembly, in *Xenopus* egg extracts, microtubule stabilization has been shown to irreversibly inhibit the formation of an NE containing NPCs^27^. Accordingly, we found that nocodazole and Aurora B inhibition had a remarkably similar effect on non-core NE assembly: Early anaphase microtubule disassembly by nocodazole reversed the non-core assembly defect on lagging chromosomes whereas there was minimal effect if nocodazole was added later (Extended Data Fig. 6b, c). Finally, during normal NE assembly, non-core NE is typically excluded from the region of the chromosome mass adjacent to dense microtubule bundles from the central spindle^9,28^. We found that this exclusion occurs whether Aurora B localizes to the spindle midzone or is forced to remain on the main chromosome mass (Extended Data Fig. 6d).

Although consistent with the hypothesis that microtubules inhibit non-core NE assembly, the above experiments do not conclusively separate the interdependent effects of Aurora B and microtubules^26^. To make this distinction, we used paclitaxel to prevent microtubule disassembly after Aurora B inhibition (Fig. 3a) and determined whether microtubules can still exclude the non-core NE from central spindle region, in the absence of active Aurora B (Extended Data Figs. 1b, 6d). In untreated controls, core (emerin) and non-core (Nup-133) proteins become intermingled on the main chromosome mass after mitotic exit as expected. By contrast, in paclitaxel-treated cells there was a persistent non-core NE gap and/or an exaggerated core NE domain at this stage (Fig. 3a, arrowheads). Importantly, this effect of paclitaxel was completely independent of Aurora B because co-addition of paclitaxel and the Aurora B inhibitor had the identical effect as paclitaxel treatment alone (Fig. 3a). Thus, microtubules inhibit non-core NE assembly irrespective of Aurora B activity.

We next asked whether the inhibitory effect of microtubules on non-core NE assembly was local and depended on the degree of microtubule bundling near the lagging chromosome. To address this, we “loosened” central spindle microtubule bundling by siRNA-mediated knockdown of the kinesin KIF4A^29^. As expected, KIF4A knockdown preserved general spindle organization and central spindle localization of Aurora B (Extended Data Fig. 7a)^29^. Consistent with our hypothesis, KIF4A depletion increased the recruitment of non-core NE to lagging chromosomes (Fig. 3b-d, Extended Data Fig. 7b, c). Interestingly, the NE at these sites often displayed a small-scale separation of core (emerin, white arrowheads) and non-core proteins (Nup133, white arrows in Fig. 3d). These “mini” core/non-core subdomains formed at any location within the central spindle, including the spindle midzone where Aurora B concentrates. Structured illumination microscopy (SIM) of the lagging chromosomes from KIF4A-deleted cells revealed that the regions of microtubule-chromosome contact, marked by the ESCRT-III component CHM4B^24^, were depleted for NPCs (Nup133) (Fig. 3e, Extended Data Fig. 8). Together, these experiments suggest that NE subdomain assembly is primarily controlled by the organization of microtubules, independent of chromosome position within the spindle.

A direct prediction of our microtubule-inhibition model is that positioning of chromosomes away from the central spindle should normalize NE assembly and restore nuclear function. To achieve a peripheral localization of missegregated chromosomes, cells were exposed to a high concentration of a MPS1 inhibitor, causing chromosome missegregation prior to the completion of chromosome congression. In HeLa cells, MPS1 inhibition was combined with the knockdown of tubulin tyrosine ligase (TTL), which further enhances peripheral chromosome localization as previously reported (Fig. 4a, Extended Data Fig. 9a)^30^. High temporal resolution, live-cell imaging was used to identify chromosomes that either remained consistently peripheral or consistently within the central spindle from anaphase onset until mitotic exit (Fig. 4a). Unlike lagging chromosomes within the spindle, the peripheral chromosomes recruited both core and non-core NE proteins, including NPCs (Fig. 4b, c, Extended Data Fig. 9a, Supplementary Video 4). As expected, microtubule and ESCRT-labeling^24^ suggested that peripheral chromosomes had less contact with microtubules than central spindle lagging chromosomes (Extended Data Fig. 9b). Strikingly, using a combination of live and fixed cell imaging, we observed that micronuclei derived from peripheral chromosomes had normal levels of nuclear import, a normal accumulation of RPA2 and lamin B1, and a normal extent of DNA replication (Fig. 4d and Extended Data Fig. 9c-e). The restoration of NE assembly and function on peripheral micronuclei, which occurred independent of their chromosome number, likely contributes to their relatively larger size (Extended Data Fig. 9c-h).

We next determined whether micronuclei from peripheral chromosomes have a lower frequency of spontaneous NE disruption and a consequently lower frequency of DNA damage (Fig. 4e). In HeLa cells, peripheral micronuclei indeed had a lower frequency of NE disruption, verified with two independent assays, and a significantly lower frequency of DNA damage (Fig. 4e, f, Extended Data Fig. 10a). In RPE-1 cells, peripheral chromosome localization also significantly reduced NE disruption and DNA damage (Extended Data Fig. 10b, Supplementary Video 5). However, in RPE-1 cells, the effect of peripheral chromosomes on NE disruption was partly masked because the large micronuclei from peripheral chromosomes in these cells are subject to actin-dependent NE breakage, as can occur transiently on primary nuclei^14^ (see discussion in Extended Data Fig. 10b, c). Thus, positioning chromosomes away from the spindle restores NE assembly and function of the resulting micronuclei.

Here, we have uncovered the basis of NE defects in micronuclei. The findings define an important mechanism underlying chromothripsis and suggest a new model for the coordination of chromosome segregation and nuclear envelope assembly during normal cell division (Supplementary Video 6). We propose that in telophase, densely bundled spindle microtubules inhibit the recruitment of non-core NE to lagging chromosomes, with the consequence that lagging chromosomes become enclosed by an NE primarily composed of core proteins. Based on prior in vitro studies^22,27^, core-only NE assembly should be a nearly irreversible barrier for the assembly of NE containing NPCs and other non-core proteins. Because micronuclei are generated with a largely-isolated core NE, they have little or no access to the nuclear transport-dependent interphase pathway for NPC assembly^12^. Defective nucleocytoplasmic transport then leads to defects in the accumulation of numerous proteins, including those necessary for normal DNA replication, DNA repair, and the maintenance of NE integrity. The resulting spontaneous NE disruption then leads to DNA damage, chromosome fragmentation, and ultimately chromothripsis^2,3,7^. Why microtubules inhibit non-core NE assembly remains to be determined, but could occur through a simple physical barrier effect. Non-core NE, including NPCs, was recently reported to assemble with fenestrated ER sheets^31^ that might less readily penetrate dense bundles of spindle microtubules than vesicles or tubules, which we speculate could be the main source of core membrane proteins.

Together, our findings demonstrate that altered NE assembly on lagging chromosomes is not the consequence of a beneficial checkpoint delay, but rather a pathological outcome. Consequently, during normal cell division, instead of precise and continuous monitoring of chromosome position, it appears that there is only loose coordination, with normal non-core NE assembly being dependent on timely spindle microtubule disassembly. Loose coordination, coupled with the irreversibility of NE assembly errors during mitotic exit, provides one explanation of why chromothripsis is common, with frequencies recently reported to be as high as 65% in some cancers^5,6^.

## Acknowledgements

We thank I. Cheeseman, T. Rapoport, N. Umbreit, T. Walther, and K. Xie for comments on the manuscript; E. Jackson and A. Spektor for preliminary experiments; J. Ellenberg, D. Gerlich, E. Hatch, M. Hetzer, A. Hyman, and T. Kuroda for reagents; J. Waters and T. Lambert of the Nikon Imaging Center at Harvard Medical School for advice and use of microscopes; M. Cicconet and C. Yapp from the Image and Data Analysis Core at Harvard Medical School for assistance with 3D rendering; Lin Shao (Yale University) for SIM reconstruction code. We acknowledge the use of the Wadsworth Center’s Electron Microscopy Core Facility. A. K. is supported by the NIH grant GM059363. D.P. is a HHMI investigator and is supported by R37 GM61345-14.

## Author Contributions

D.P., S.L., and M.K. designed the experiments. D.P., S.L., and M.K. wrote the manuscript, with edits from all authors. S.L., and M.K. performed most experiments and analysis. M.M. assisted with several experiments and contributed Extended Data Fig. 10b, c. N. Y. and A. K. performed the electron microscopy.

